# The co-evolution of adhesion and F-actin gives rise to hierarchical cellular force generation

**DOI:** 10.64898/2026.11.08.765126

**Authors:** Yuanning Guo, Noam Ottolenghi, Malak Amer, Jiayu He, Eran Bouchbinder, Haguy Wolfenson

## Abstract

Cells — the basic units of life — dynamically adhere to their micro-environment, build their F-actin cytoskeleton, and apply molecular-motor generated contractile forces to it in a remarkably broad range of biological processes and physiological functions, in both health and disease. Yet, how these processes are orchestrated over multiple spatiotemporal scales is not fully understood. Using several experimental approaches and a theoretical framework, we show that adhesion assembly, on-adhesion F-actin accumulation and the contractile force itself all follow the same time evolution over tens of minutes, the cellular force timescale, at well-defined adhesion sites. We also show that the F-actin distribution away from the adhesion site features a multiplicative space-time decomposition. Finally, we show that F-actin retrograde flow — central in the so-called molecular motor-clutch models — is markedly faster than force generation rates, that it does not slow down over maturing adhesions and that its density drops at adhesion sites. These findings suggest a hierarchical, multi-scale force generation process that is controlled by slow adhesion recruitment, which is accompanied by rapid, bursty linear actin polymerization. The linear F-actin slowly accumulates with adhesion recruitment to form an extended near-adhesion F-actin network to which molecular motors attach and apply elastic-like contraction.

---

Cells possess remarkable abilities to adhere to their surroundings, dynamically remodel their cytoskeleton, and actively apply contractile forces to them. Adhesion and force generation fundamentally affect a broad range of processes involving mechanical control of cellular motion, shape, signaling and fate. These include cell spreading and polarization [1–3], cell migration [4– 6], mechanosensing and mechanotransduction [7–9], cell differentiation [10, 11], proliferation [12, 13], and apoptosis [14, 15], among other fate decisions. As such, cellular adhesion and force generation are essential for numerous physiological functions [16–18]. How do cells regulate their adherence to the environment and the application of force to it? How are these basic intracellular and extracellular processes coordinated over space and time?

The basic molecular building blocks involved in adhesion and force generation are mostly known. Adhesion is realized through molecular linkages (“clutches”) [19, 20], composed of transmembrane integrins and related adaptor proteins, that mechanically connect the dynamic cell cytoskeleton to its environment, the extracellular matrix (ECM). On the extracellular side, the integrins attach to ligands, i.e., ECM proteins such as fibronectin and collagen. On the intracellular side, the integrins are connected via adaptor proteins (e.g., talin, vinculin) to an actomyosin network, composed of filamentous actin (Factin) that undergoes continuous polymerization and depolymerization and to which myosin II molecular motors attach. These motors convert chemical energy into sliding motion of the actin filaments (through ATP hydrolysis), giving rise to contractile forces that are transmitted to the ECM through the adhesion complexes [21].

The ways in which these molecular building blocks collectively give rise to adhesion and force generation have been subjected to extensive research, both experimental and modeling oriented [19, 20, 22–38]. These efforts led to the molecular motor-clutch class of models [20, 29, 31, 33, 35–38], which typically consider a single actin filament that moves from the cell edge inward at a velocity that is determined by the model. Traction forces are transmitted from this retrograde actin flow to the ECM in a friction-like manner through the stochastic engagement of linkages, involving binding and unbinding kinetics, both to the moving actin filament and to the ECM. This picture of cellular adhesion and force generation usually lacks spatial aspects and is confined to a timescale of seconds or less, significantly shorter than the timescales associated with the aforementioned cell-scale processes.

Far less is known and understood about how these multiple building blocks mutually interact and self-organize in space and time at intermediate spatiotemporal scales, i.e., at and near adhesion sites during their lifetime of the order of tens of minutes. Addressing this fundamental problem involves various challenges. First, the dynamics of individual adhesions alone span a broad range of scales, involving timescales ranging from a fraction of a second to tens of minutes and spatial scales ranging from nanometers to several microns [39–42]. Second, quantifying the spatiotemporal co-evolution of various adhesion, cytoskeletal and force generation processes — which is a major challenge on its own — requires to cleanly isolate adhesions from birth to disassembly, and to precisely identify the actomyosin dynamics and contractile force that are associated with a particular adhesion, without the interference of nearby, coexisting adhesions. Third, the generality of any findings should be tested/demonstrated across various adhesion landscapes, cell types and cellular states (e.g., spreading and migratory).

## Experimental approach

To address these challenges, we employed three experimental approaches, the first of which involves cells adhering to hexagonal arrays of ligand-coated, flexible micropillars (Fig. 1A). Under these conditions, adhesions can be formed on a well-defined “adhesion landscape”, i.e., exclusively at the pillars’ top surface, which are placed at a priori known and sufficiently well separated positions. This allows to cleanly identify “isolated adhesion events” occurring near the cell edge (Supplementary Text, sect. S-1), without the potential interference of nearby, coexisting adhesions. The adhesion events we consider hereafter all correspond to maturing adhesions (from their initial assembly), which are significantly larger and longer-lived compared to nascent, non-maturing adhesions [43]. In parallel, the time-dependent generated force is accurately probed by measuring the pillar displacement/deflection Δ(*t*), using time-lapse microscopy (Fig. 1A and Methods). We use Δ(*t*) instead of the force *F* (*t*) (obtained by multiplying the former by the pillar’s flexural rigidity) as there is evidence that it is the quantity controlled by cells [9].

**Fig 1.**
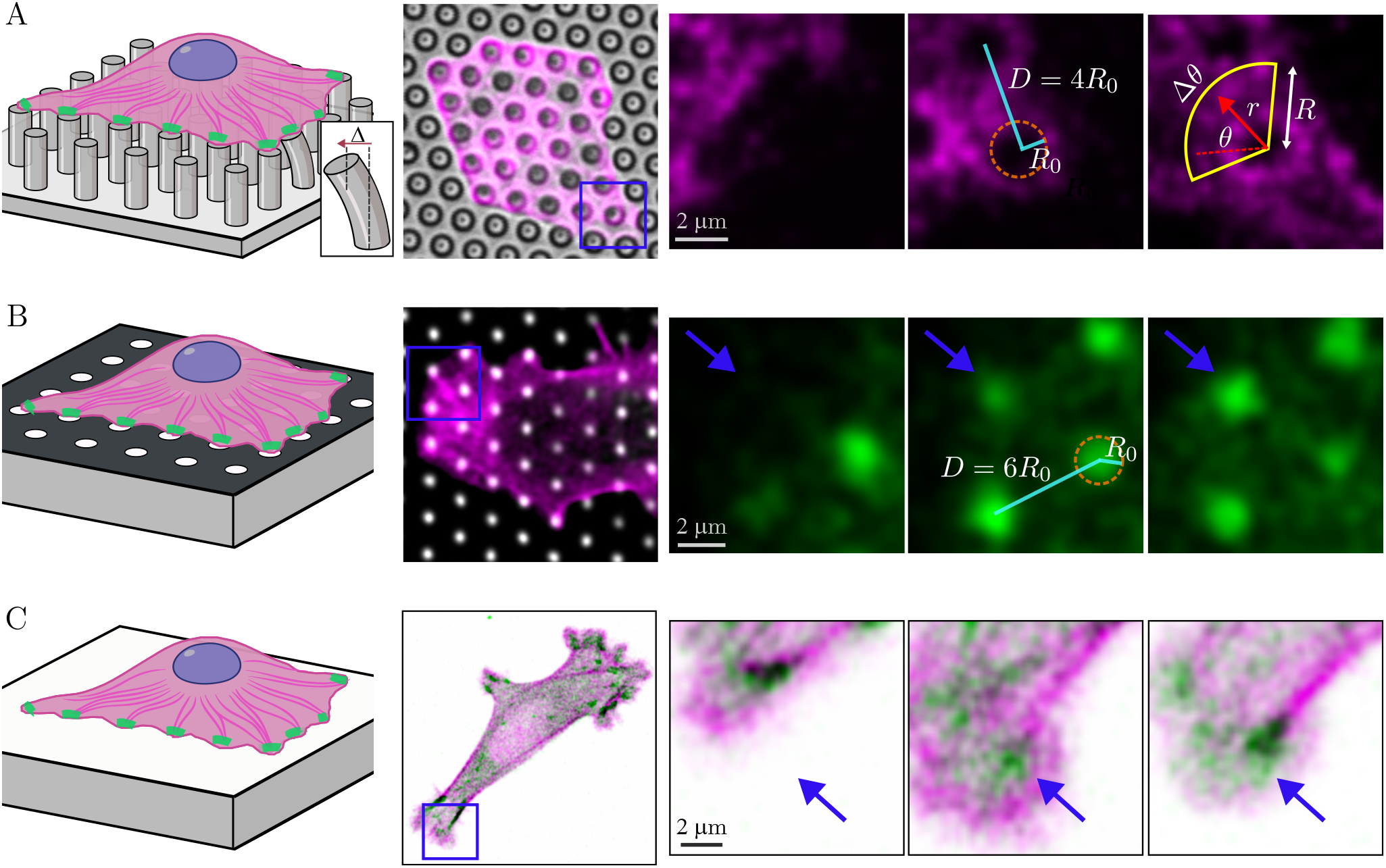
The three experimental approaches. (**A**) A fibronectin-coated micropillar array system, (**B**) a flat substrate micropatterned with a hexagonal array of circular fibronectin islands (white; black areas are passivated regions without fibronectin), (**C**) a continuously fibronectin-covered substrate. Representative frames from live-cell imaging experiments are presented for each system, showing cells coexpressing tdTomato-Tractin (magenta, for F-actin) and GFP-paxillin (green, for adhesion). The three images on the right of each panel are magnified views of the regions outlined by the blue squares, showing the evolution of a near-adhesion F-actin network in (A) and (C), and/or adhesion assembly and maturation in (B) and (C) (blue arrows). The dashed circles in (A) and (B) indicate the pillar top surface and fibronectin island outlines, respectively, with *R*_0_ depicting the radius (1 *µ*m in both cases) and *D* the center-to-center distance. Pillar deflection, Δ(*t*), is used for measuring traction forces, and Δ*θ*, is the angular sector used for coarse-graining various radii, *R*, are shown in (A). Scale bars, 2 *µ*m. See text, Methods and Supplementary Text for additional details.

We also simultaneously measure the spatiotemporal evolution of both the amounts of adhesion and F-actin that are uniquely associated with the generated force. This is achieved by high-resolution fluorescence microscopy (Methods), involving a GFP-labeled adhesion protein whose measured intensity gives rise to a density field *a*(*r, θ, t*) (Fig. 1A, here *r* and *θ* are polar coordinates located at the center of each relevant pillar), and fluorescently-tagged F-actin, whose measured fluorescent intensity gives rise to a density field *c*(*r, θ, t*) (Fig. 1). Finally, by refining the measurement resolution, we extract the actin flow speed *v*(*t*) along selected lines (see below). Later on, two additional, complementary experimental approaches will be discussed.

### The spatiotemporal dynamics of cellular force generation

The first question we address concerns the relation between the adhesion-level time evolution of the amounts of F-actin and adhesion. To that aim, we define the coarsegrained quantities 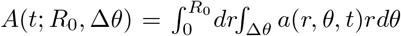 and 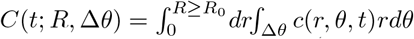, where *R*_0_ = 1 *µ*m is the radius of the pillar (i.e., the region where adhesion can possibly form). Δ*θ* is an angular sector generally pointing to the proximal part of the adhesion (Fig. 1A) of a sizable fraction of 360^*°*^ (2*π* radians), whose physical significance will be discussed below. Note that in *C*(*t*; *R*, Δ*θ*) one can have *R ≥ R*_0_ since the F-actin network extends also away from the adhesion. In Fig. 2A, we superpose *A*(*t*; *R*_0_, Δ*θ*) and *C*(*t*; *R*_0_, Δ*θ*), each with a unity amplitude, for an adhesion event in a human breast adenocarcinoma cell (MDA-MB-231).

**Fig 2.**
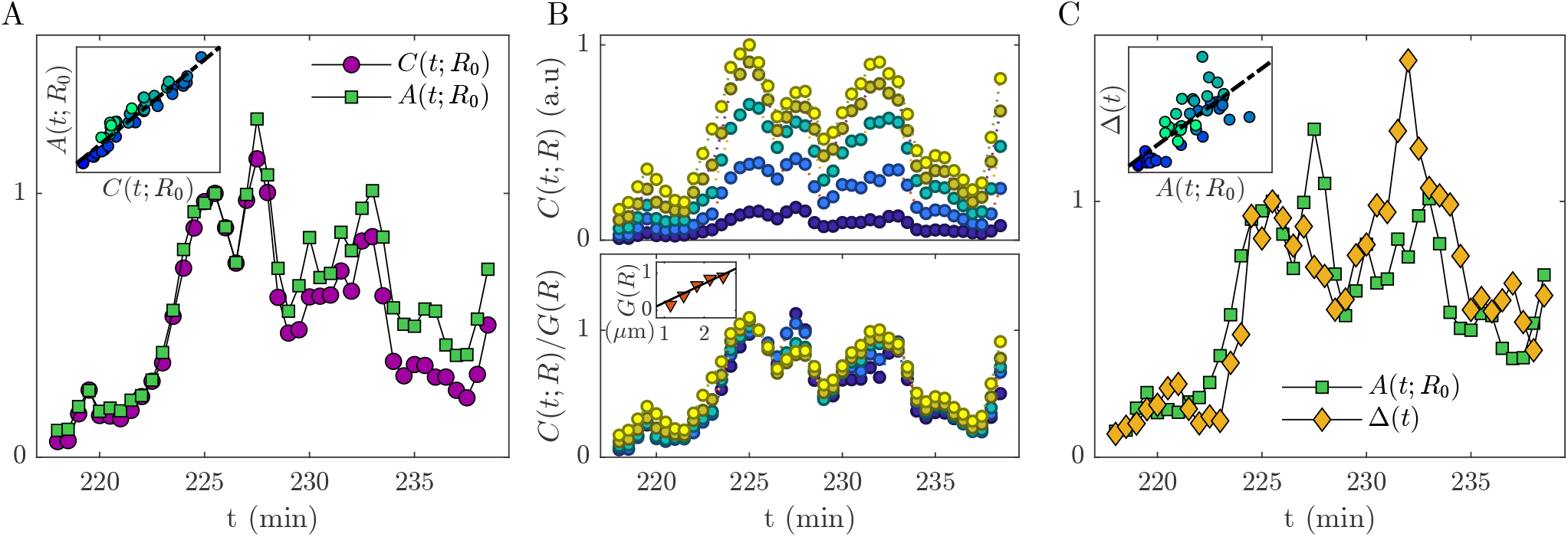
The spatiotemporal dynamics and co-evolution of adhesion assembly/disassembly, F-actin density and adhesion-level generated force during adhesion’s lifetime. (**A**) The integrated amount of adhesion *A*(*t*; *R*_0_, Δ*θ*) and of on-adhesion F-actin *C*(*t*; *R*_0_, Δ*θ*) (both normalized to unity amplitude), see legend and text for definitions, for an isolated adhesion event in a quasi-stationary spreading MDA-MB-231 cell adhering to a micropillar array. The pillars feature a fibronectin-coated top surface of radius *R*_0_ = 1 *µ*m and a center-to-center distance of *D* = 4*R*_0_. The integration angular sector is Δ*θ* = 240^*°*^ (see text and Fig. 1A and Supplementary Text, and note that it is omitted in the legend expressions for brevity). To avoid spatial discretization effects we actually set *R* = 1.16*R*_0_ in the integration domain (Supplementary Text, sect. S-2). The results support Eq. (1), see also the strong time correlation in the inset (time increases from blue to green). (**B**) (top) The time evolution of *C*(*t*; *R*, Δ*θ*) away from the adhesion for (from bottom to top) *R/R*_0_ = 1.16, 1.49, 1.82, 2.15, 2.48. (bottom) The same as the top panel but for *C*(*t*; *R*, Δ*θ*)*/G*(*R*), where the amplitude *G*(*R*) *∼ R* is presented in the inset. The results support Eq. (2). (**C**) The same as (A), but for the normalized adhesion-level contractile displacement Δ(*t*) (proportional to the force *F* (*t*)) instead of *C*(*t*; *R*_0_, Δ*θ*), see legend. The results support Eq. (3). See also movie S1.

The two quantities are observed to follow the very same time dependence over the entire adhesion event duration (here *∼* 20 min [24]), from adhesion onset to disassembly. This co-evolution (see also inset in Fig. 2A) persists down to short-time features (e.g., an oscillatory behavior on a scale of *∼* 4 min [9, 44, 45]) and is found for many other adhesion events (Supporting Figures and Supplementary Text), implying that

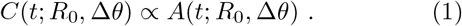

It suggests that each small adhesion patch — composed of integrins and adaptor proteins — is connected to a fixed amount of F-actin at any time, a relation that is not simply inferred from a bottom-up molecular picture [46, 47]. While the adhesion is localized on a scale *R*_0_, the F-actin network extends further away towards the cell body, featuring *R > R*_0_. In Fig. 2B(top), we present five *C*(*t*; *R*, Δ*θ*) curves for *R*_0_ *<R<* 2.5*R*_0_ (from bottom to top, see caption), for the adhesion event shown in Fig. 2A. In Fig. 2B(bottom), we plot the very same curves, each normalized by its amplitude *G*(*R*) (defined at the same time point for all curves and shown in the inset). Remarkably, all curves almost perfectly collapse on a master curve *T* (*t*), implying

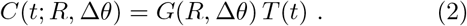

This multiplicative space-time decomposition is found for many other adhesion events (Supporting Figures and Supplementary Text), indicating *c*(*r, θ, t*) *∼g*(*r*)*T* (*t*) with 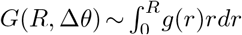, to a good approximation. These findings imply that the spatiotemporal formation of the F-actin network behind the adhesion is such that the F-actin density in each small region in space approximately follows the same time evolution. Moreover, we find *G*(*R*) *∼ R* to a good approximation (inset of Fig. 2B(bottom)), implying a decaying *g*(*r*) *∼* 1*/r* density profile (Supplementary Text, sect. S-3).

We next consider the contractile force, quantified through Δ(*t*). It has been shown in [9] that Δ(*t*) *∝ C*(*t*; *R*, Δ*θ* = 2*π*) for a fixed *R>R*_0_, which together with Eqs. (1)-(2), implies

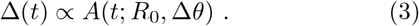

This prediction is verified in Fig. 2C and is similarly demonstrated for many other adhesion events (Supporting Figures and Supplementary Text). A related observation has been made in [25], but only during the initial stages of adhesion maturation and force buildup. Equation (3) also clarifies the role of Δ*θ* in both *A*(*t*; *R*_0_, Δ*θ*) and *C*(*t*; *R*, Δ*θ*) in Eqs. (1)-(2). That is, we are interested in the part of the F-actin network that can contribute to contractile force generation and thus focus on a wide angular sector Δ*θ* pointing towards the cell body and likewise use it in *A*(*t*; *R*_0_, Δ*θ*) (Supplementary Text, sect. S-2).

The results of Fig. 2 (see also movie S1, along with the robustness of the findings across many other adhesion events, Supplementary Text) gives rise to the following picture of cellular force generation at maturing adhesions: the amount of adhesion and the generated force follow the same time evolution over the adhesion’s lifetime on the scale of *∼* 10 min, whereas the F-actin density at and behind the adhesion approximately follows the very same time evolution at each spatial patch. It suggests that adhesion accumulation/maturation is the key underlying process that controls actin polymerization and the formation of an F-actin network behind the adhesion, which in turn provides available sites to which myosin II motors attach and generate a spring-like displacement [9, 26].

Moreover, the above picture suggests that linear, quasi-one-dimensional actin filaments are polymerized radially from small adhesion patches, which is perfectly consistent with a spatial coarse-grained profile *g*(*r*) *∼* 1*/r*, as we generically observe (Supplementary Text, sect. S-3). This actin polymerization process is likely to be very fast (possibly subsecond [48]), yet it is the slow, minutes-scale adhesion accumulation process, characterized by the temporal function *T* (*t*), which controls force generation. The formation of quasi-one-dimensional radial actin filaments gives rise to an extended F-actin network that results in the space-time decomposition in Eq. (2). This hierarchical spatiotemporal picture also raises questions regarding the role of retrograde actin flow in force generation, to be discussed below.

### The broader generality of the results

To further support the generality of our observations, we discuss two additional experimental approaches. The micropillars approach introduces in addition to an adhesion/ligand landscape also a non-planar ECM topography. Hence, we also employed a complementary approach that involves micropatterning of planar, continuous substrates. The latter are micropatterned with a desired pattern of ligand islands (Fig. 1B), retaining the ability to cleanly identify isolated adhesion events, with the added benefit of flexibly controlling the ligand islands size, shape and separation. At the same time, it does not allow to directly measure the force itself.

In Fig. 3(top row), we present results for a micropatterned ECM with a hexagonal array of circular ligand islands of same radius *R*_0_ = 1 *µ*m as in the micropillars experiments in Fig. 2. To further test for generality, we made three additional changes compared to the latter experiments: (i) we increased the center-to-center distance between the ligand islands from 4*R*_0_ to 6*R*_0_, (ii) we considered a different cell type, a mouse embryonic fibroblast (MEF), (iii) we analyzed a different cellular state, a migratory state instead of a quasi-stationary spreading state. The results in panels (A)-(C) strongly support the validity of Eqs. (1)-(2), despite the significant changes made in all above-discussed respects. While the generated force was not measured, the validity of Eqs. (1)-(2) implies the validity of Eq. (3) as well. In panel (D), we present both the time evolution of the cell area and its center-of-mass displacement, demonstrating the migratory cellular state. The results in Fig. 3(top row) are validated for many other adhesion events (Supporting Figures and Supplementary Text).

**Fig 3.**
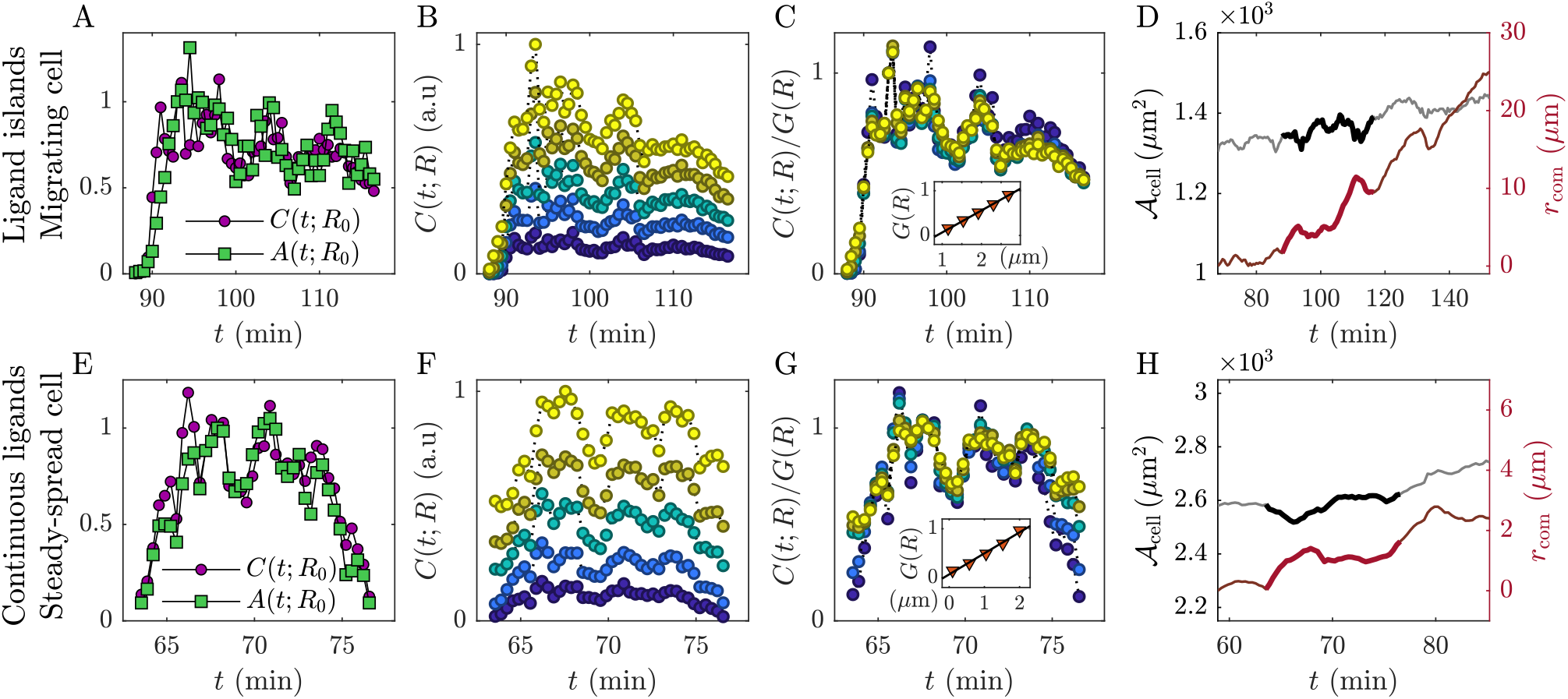
The generality of the findings. (**A**)-(**C**) The same as Fig. 2A-B but for an isolated adhesion event in a MEF migrating on a planar, continuous substrate with micropatterned fibronectin. The latter feature a hexagonal pattern of circular ligand islands (Fig. 1B) of radius *R*_0_ = 1 *µ*m and a center-to-center distance of *D* = 6*R*_0_. The results support Eqs. (1)-(2) with *G*(*R*) *∼ R*, see text for discussion and Supplementary Text for additional technical details. (**D**) The migratory cellular state is demonstrated by plotting the cell’s area *A*_cell_(*t*) (left *y*-axis), which is quasi-stationary, and its center-of-mass position/distance *r*_com_(*t*) (right *y*-axis), which significantly evolves over the time window analyzed in A-C (highlighted in a thicker line). (**E**)-(**H**) The same as A-D but for an isolated adhesion event in a quasi-stationary spreading MEF adhering to a planar, continuous substrate with a continuous fibronectin coverage (Fig. 1C). The quasi-stationary spreading cellular state is demonstrated in *H*, where both *A*_cell_(*t*) and *r*_com_(*t*) vary mildly over the time window of interest (again highlighted in a thicker line). The results support Eqs. (1)-(2) with *G*(*R*) *∼ R*, see text for discussion and Supplementary Text for additional technical details.

Finally, we employed a third experimental approach involving a continuous ligand coverage of planar, continuous substrates (Fig. 1C). Here, adhesions can form anywhere and the identification of isolated adhesion events is more challenging, yet not impossible. Inspired by the analysis above, this has been achieved (Supplementary Text) and indeed the generality of our results is further demonstrated in Fig. 3(bottom row) for a nearly stationary spreading cellular state. Additional supporting examples can be found in the Supporting Figures and Supplementary Text.

### Assessing the role of retrograde actin flow in cellular force generation

As mentioned above, retrograde actin flow plays a central role in clutch models of cellular force generation/transmission. In this framework, F-actin flows from the cell edge towards its center, at a speed *v* that is self-consistently selected by a given model. The interaction between this retrograde flow and the ECM through molecular linkages gives rise to contractile forces. Specifically, in the “strong adhesion” regime that is relevant for our measurements, molecular linkages between the flowing actin and the ECM are formed at a high rate, leading to the slowing down of the flow (i.e., to a reduction in *v*) that is accompanied by friction-like force *F* buildup, e.g., [20, 22, 23, 33–38]. Notably, these models typically do not feature spatial aspects, do not account for the evolution of the F-actin network and their dynamics are fast, i.e., reach a stationary friction-like force-velocity *F* (*v*) relation on the scale of seconds [38]. Here, we shed light on the role of actin flow in force generation in view of our findings so far.

For the dense and uniform fluorescent labeling used above to measure *c*(*r, θ, t*), information about *v* is extracted by focusing on a region in space, e.g., a line going through the adhesion (Fig. 4D-E), and using *c*(*r, θ, t*) along it to generate a kymograph, as explained next. To this aim, we performed the same experiments as above with increased spatiotemporal resolution (of *∼* 120 nm, using 40 nm/pixel, and 1.6 sec, respectively). In Fig. 4A(top), we present such a kymograph for an adhesion event on a micropatterned substrate (movie S2), where we also superpose the corresponding results for *a*(*r, θ, t*), clearly revealing adhesion assembly. *v* can be then extracted only if the actual flow is intermittent, i.e., characterized by ‘bursts’ manifested as diagonal streaks in the F-actin kymograph (whose slope corresponds to *v*). Such kymograph features are indeed revealed when the adhesion protein and F-actin channels are separated, see the two subsequent sub-panels.

**Fig 4.**
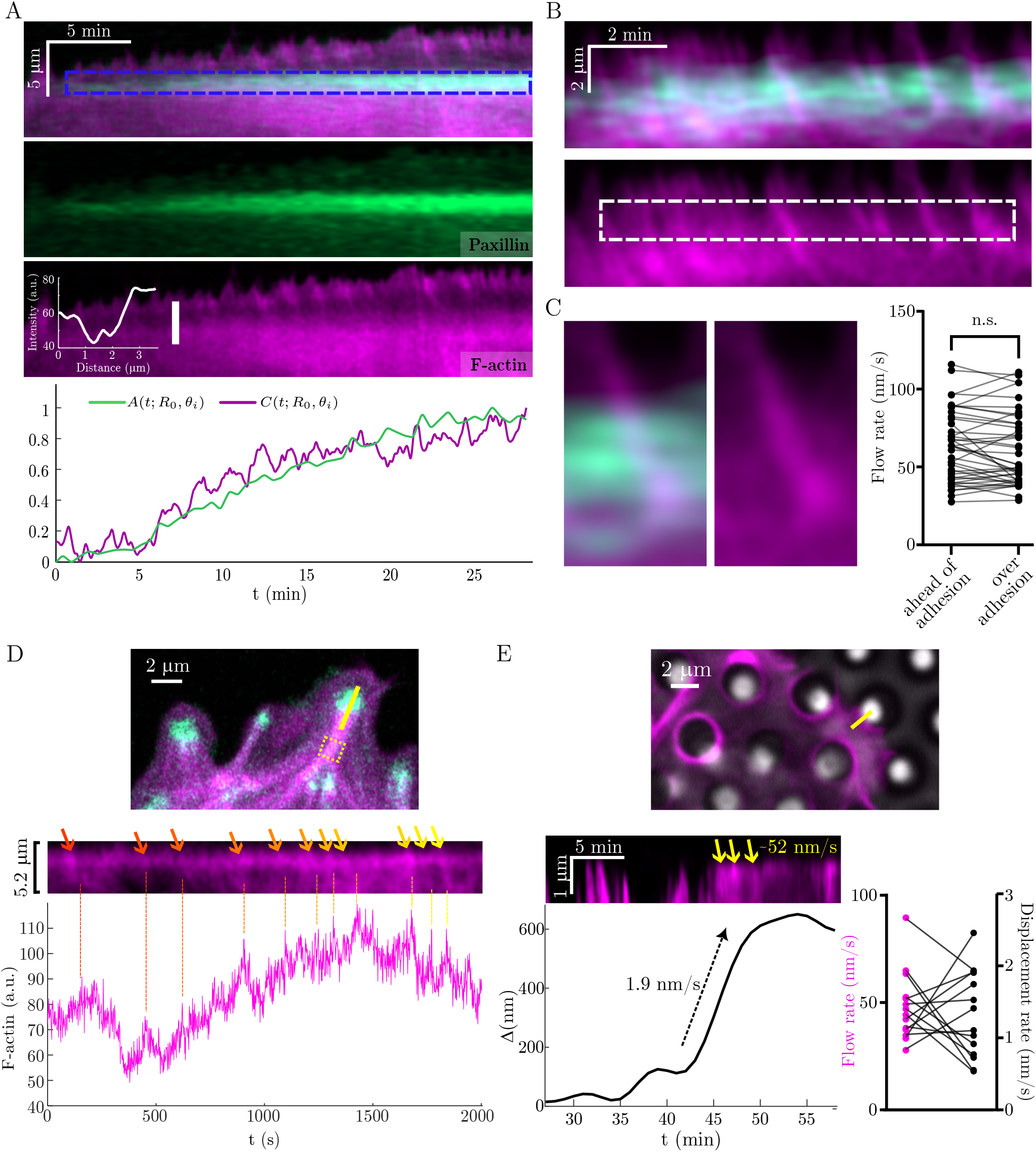
Retrograde F-actin flow during force generation. (**A**)-(**B**) Kymographs generated for adhesion events on micropatterned substrates, each along a line going through the adhesion at an angle *θ*_*i*_, in cells co-expressing tdTomato-Tractin (magenta, for F-actin) and GFP-Paxillin (green, for adhesion). The blue rectangle in (A) denotes the adhesion of linear size *R*_0_. The inset in the lower kymograph in (A) shows the intensity profile along the white line. The coarse-grained intensities *A*(*t*; *R*_0_, *θ*_*i*_) and *C*(*t*; *R*_0_, *θ*_*i*_) are plotted at the bottom of (A) (Supplementary Text, sect. S-2, and see also movie S2). Diagonal streaks in (B), corresponding to actin polymerization bursts, traverse the adhesion (outlined by the white rectangle) without significant change in slope. (**C**) Higher-magnification of a single actin polymerization burst traversing an adhesion (left) and a quantification of actin flow rates ahead of an adhesion vs. over the corresponding adhesion for multiple bursts (right). (**D**) Frame from a live-cell imaging experiment (same color code as above), where the yellow line was used to generate the kymograph. The yellow square marks the area used to quantify F-actin (bottom). (**E**) Similarly to (D) but in a micropillar array experiment. Actin flow rates and contractile displacement rates 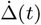 are compared (bottom-left for the kymograph and bottom-right for multiple pillars).

The F-actin kymograph also clearly reveals the accumulation of F-actin behind the adhesion, extensively discussed in Figs. 2-3. As noted above, while Eqs. (1)-(2) involve areal integration, we expect them to be valid for the underlying densities and consequently also for lines going through the center of the adhesion. Indeed, performing such line integrals gives rise to the excellent agreement between the time-dependent amount of adhesion and F-actin shown in Fig. 4A(bottom), further strengthening the results in Figs. 2A and 3A,E. A closer inspection of the F-actin kymograph reveals that the density of F-actin significantly drops at the adhesion site, compared to its values both ahead and behind it, see the superposed graph. This has been observed in the majority of cases evaluated (24/32 kymographs). Furthermore, despite the density reduction, the actin speed does not appear to drop.

To highlight the latter, we focus on another adhesion event in Fig. 4B in which the drop in F-actin density over the adhesion is not as pronounced as in Fig. 4A. Multiple polymerization bursts that traverse the adhesion are evident (see also further zoom-in on one such event in Fig. 4C(left)). Quantification of multiple such events from multiple adhesions comparing flow rates ahead of the adhesion vs. over the adhesion clearly reveal that the flow does not slow down at the adhesion (Fig. 4C(right)). This observation does not support a dynamic friction-like force generation as a direct consequence of F-actin flow [32]. Instead, following the above-discussed picture, the flow appears to contribute to the buildup of the F-actin network behind the adhesion, which in turn leads to an elastic-like (as opposed to friction-like), motor-driven force generation over longer timescales. Indeed, in Fig. 4D we show that bursts of actin flow in the vicinity of an adhesion are correlated with an increase in the actin density further behind the adhesion, supporting the idea that F-actin transport from the adhesion vicinity contributes to the accumulation of F-actin behind it. Finally, results presented in Fig. 4E for an adhesion event in a micropillars experiment show that the F-actin flow is much faster than the adhesion-scale displacement rate 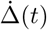, in line with the accumulation of fast and intermittent, low-dimensional adhesion-actin processes that collectively give rise to elastic-like force generation over slower timescales.

## Discussion

We elucidated the spatiotemporal processes underlying cellular adhesion and force generation, revealed through the quantitative relations in Eqs. (1)-(3) (with *G*(*R*) *∼R*) and in Fig. 4 between the basic biophysical observables involved. These include the spatiotemporal distributions of adhesion and F-actin, the F-actin flow speed and the time evolution of the adhesion-scale generated force. Our findings show that force generation crucially depends on the spatiotemporal co-evolution of adhesion and its surrounding F-actin network, where the latter is an essential, inseparable part of the process. The multiplicative space-time property of the F-actin distribution indicates that the slow, minutes-scale force buildup emerges from coarse-graining over low-dimensional, fast and intermittent adhesion-actin processes, overall giving rise to a hierarchical, multiscale force generation. Myosin II motors activity depends on the formation of the F-actin network, and hence is ‘slaved’ to its slow timescale, and the motor-generated contractions give rise to a spring-like force, rather than to a dynamic friction-like force that depends on the actin flow rate.

This hierarchical, multiscale picture of cellular force generation is not captured by existing models. Specifically, clutch models may pertain to the low-dimensional and fast processes, but require extensions and coarsegraining to be relevant for adhesion-scale force generation, necessarily taking into account the spatiotemporal co-evolution of adhesion and the F-actin network. Yet, we do not find strong evidence for a dynamic friction-like force generation that directly emerges from the retrograde actin flow assumed in these models. Specifically, we find that F-actin flow does not slow down over maturing adhesions and its density drops therein, findings that appear to be consistent with the speckle microscopy measurements of [27]. Instead, bursts of F-actin flow feed in to the slow buildup of the F-actin network required for elastic-like force generation.

Our results raise various interesting questions for future investigation. First, one should understand the origin of the characteristic timescale of adhesion and force generation, of the order of 10 minutes [24]. It is a basic cellular timescale, with potential implications for many other cellular processes, including integration of signals from multiple adhesion sites toward cell fate decisions [15]. Second, it would be interesting to explore possible differences between healthy and diseased cells and between young and aging cells in relation to adhesion and force generation. Finally, the latter are essential for mechanosensitivity, i.e., the dependence of cellular responses on the mechanical properties of the ECM [7– 9, 26, 49, 50]. It has been recently established that the contractile displacement Δ(*t*) is independent of the ECM rigidity for maturing adhesions [9]. Combining this fundamental property with Eqs. (1)-(3) gives rise to interesting predictions that should be tested in future work by systematically varying the ECM rigidity. In addition, adhesions are known to fail to mature on sufficiently soft ECMs [4], which is a regime that should be also studied in subsequent work.

## Methods

### Cell culture and fluorescent reporters

MEFs stably expressing paxillin-EGFP were provided by Benny Geiger (Weizmann Institute of Science), and MDA-MB-231 cells were provided by Yuval Shaked (Technion). Cells were maintained at 37^*°*^C and 5% CO_2_ in Dulbecco’s modified Eagle medium supplemented with 10% fetal bovine serum and penicillin-streptomycin (100 IU/mL; Biological Industries). For dual-color imaging, paxillin-EGFP MEFs were transfected with a tdTomato-Tractin construct using jetOPTIMUS (Polyplus), following the manufacturer’s instructions, and imaged *∼* 48 h later. pCMV-tdTomato-Tractin plasmid was provided by the late Mike Sheetz. For dual color measurements using the MDA-MB-231 cells, tdTomato-Tractin and GFP-vinculin were transfected using Lipofectamine 3000 (Invitrogen).

### Preparation of continuous PDMS substrates

Planar substrates were prepared from two-component PDMS (DOWSIL CY 52-276) mixture using a 1:3.2 ratio of component A to component B for an elastic modulus of 40 kPa [9]. The amount of PDMS was adjusted to the substrate area to produce an approximately 50-*µ*m-thick layer on glass coverslips. After uniform coating, substrates were cured at 70^*°*^C for 2 h.

### Surface passivation and fibronectin photopatterning

Circular fibronectin islands were produced using the PRIMO photopatterning module (Alvéole), controlled through Leonardo software. This procedure uses light-induced molecular adsorption to restrict protein deposition to illuminated regions. 40 kPa PDMS surfaces were first rendered hydrophilic using a Diener electronic plasma system (ZEPTO-W6), with the device settings of pressure 0.5 mbar, 50% power for 90 s. Within 2 h of plasma treatment, substrates were coated with poly(L-lysine)-graft-poly(ethylene glycol) (PLL-g-PEG, 100*µ*g/mL, SuSoS, SZ42-90) for one hour at room temperature. The PEG layer suppressed nonspecific protein adsorption and cell adhesion outside the intended patterns. The photoactivatable reagent PLPP (Alvéole) was then applied, and selected regions were illuminated with UVA (dose 3000 mJ/mm^2^) through the PRIMO system to locally remove passivation. After illumination (and PBS washing), substrates were incubated with fibronectin at a total concentration of 10 *µ*g/mL (37^*°*^C, 1 h), of which 30% was fluorescently labeled with Alexa Fluor 647. Excess protein was rinsed away, and the fluorescent patterns were inspected before cell seeding. Fibronectin was labeled using the Alexa Fluor 647 Antibody Labeling Kit (A20186, Invitrogen), following the kit instructions. Pattern preparation and subsequent cell experiments were completed within 24-48 h. Patterns were designed in Fiji/ImageJ within a nominal PRIMO digital micromirror device field (DMD) of 500 *×* 300 *µ*m; larger patterns were divided into multiple illumination fields by Leonardo.

### Micropillar fabrication and fibronectin coating

Sylgard 184 PDMS base and curing agent (Dow Corning) were mixed at 10:1 and cast into molds containing holes of defined depth and spacing. The molds were placed onto glass-bottom 35-mm dishes (D35-20-0-N, Cellvis), and the PDMS was cured at 60^*°*^C for 12 h. After cooling, the molds were removed while sub-merged in ethanol to reduce pillar collapse. Ethanol was subsequently exchanged for PBS. The arrays had circular pillar cross-sections of 2 *µ*m diameter and 4 *µ*m center-to-center spacing. Human plasma full-length fibronectin (FC010, Merck) was applied at 10 *µ*g/mL for 1 h at 37^*°*^C for surface coating, followed by PBS washes. The pillar flexural rigidity was calculated using Euler–Bernoulli beam theory: 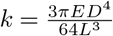, where *E* = 2 MPa is the Young’s modulus of PDMS, D is pillar diameter, and *L* = 5.3 *µ*m is pillar height.

### Cell preparation and live imaging

On the experiment day, cells were trypsinized, centrifuged in growth medium, resuspended in HBSS supplemented with 10% FBS, 20 mM HEPES, and 1% penicillin-streptomycin, and incubated at 37^*°*^C for 30 min before plating onto fibronectin-coated pillar arrays, patterned planar substrates, or uniformly coated substrates. Long time-lapse imaging was performed at 37^*°*^C using a Zeiss LSM 800 confocal microscope with an Airyscan detector with a 20 *× /*0.9 NA objective (157 nm/pixel), 488-nm excitation for GFP-paxillin, 561-nm excitation for tdTomato-Tractin, and acquisition every 30 s. For high-speed imaging of actin retrograde flow, a 63 *× /*1.4 NA oil objective was used in Airyscan mode (reaching *∼* 120 nm spatial resolution), with 3 *−* 5*×* optical magnification, reaching 40-nm pixel size, and frame intervals of 1.6 s. Tractin images were acquired in bursts of 20 consecutive frames, interleaved with GFP-paxillin imaging (every 30 s).

### Pillar tracking and displacement measurements

Pillar positions (imaged via brightfield imaging) were tracked in Fiji/ImageJ using the NanoTracking plugin, based on a cross-correlation method. Only pillars un-occupied by cells at the beginning of a recording were selected for tracking relative to an unloaded reference position. For a reference position (*x*_0_, *y*_0_), the displacement vector was obtained from the position at time t, and its magnitude was calculated as follows: 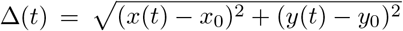. To measure and evaluate the stage drift during imaging, five randomly selected micropillars unaffected by external disturbances were used as reference markers for nano-tracking. The stage drift in both the x and y directions was calculated for each frame. Specifically, drift correction was performed by subtracting the average displacement of these five reference pillars from the measured displacement of each targeted pillar in both directions. To evaluate systemic noise, another unaffected pillar from a different location was selected; its residual displacement, obtained by subtracting the corrected drift, was defined as the noise level. All corresponding operations and data analysis were implemented in MATLAB (v2018b).

Photobleaching correction was applied selectively based on manual assessment of quantitative whole-cell fluorescence intensity time courses. Whole-cell integrated fluorescence intensity was measured in each frame and plotted against time. Sequences showing a clear, sustained overall decline consistent with photobleaching were selected for correction; no photobleaching correction was applied to sequences without an apparent decay. For selected sequences, the intensity time course was fitted to a single-exponential decay model with a constant offset, 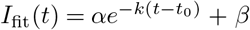, using nonlinear least-squares fitting initialized by a grid search. Here, *α* is the decay amplitude, k is the decay rate constant, *β* is the constant offset, and *t*_0_ is the time of the first analyzed frame. For image correction, the frame-specific mean background intensity measured in cell-free regions was first subtracted from pixel intensities. All pixels within each frame were then multiplied by the same correction factor, 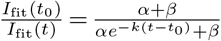, thereby referencing the correction to the fitted initial intensity. Fitting and correction were implemented using custom R (v4.4.0) and Python (v3.12.0) scripts.

## Acknowledgements

We acknowledge the assistance of the following members of the Wolfenson lab with training and experimental procedures: Elisabeth Nadjar Boger, Lea Feld, Shay Melamed and Lidan Shi. We also thank Maya Holdengreber (Technion Biomedical Core Facility) for technical assistance. This work was made possible in part through the support of grant 63526 from the John Templeton Foundation (E.B. and H.W., the opinions expressed in this publication are those of the authors and do not necessarily reflect the views of the John Templeton Foundation). H.W. also acknowledges support from the Israel Science Foundation (grant 1984/24).

## Author contributions

E.B. and H.W. conceived and designed the research, supervised the research, acquired funding and wrote the original manuscript. Y.G. performed the extensive experiments, contributed to their analysis, led the visualization of the experimental data and reviewed the manuscript. N.O. led the analysis of the experimental data, their theoretical interpretation, significantly contributed to the visualization of the experimental data and reviewed the manuscript. M.A. performed some of the experiments involving the MDA-MB-231 cells. J.H. significantly contributed to the micropillar tracking analysis. E.B., H.W., Y.G. and N.O. extensively discussed the results.

## Supporting Figures

**Supporting Fig 1.**
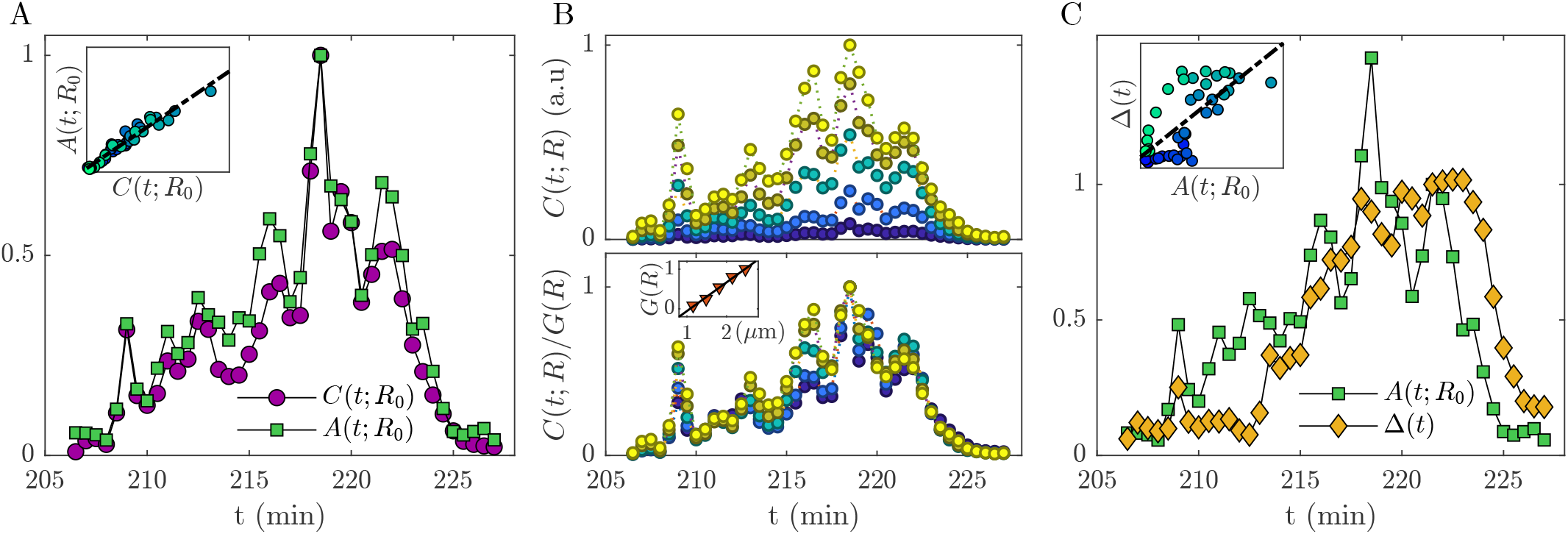
An additional example of an isolated adhesion event in a quasi-stationary spreading MDA-MB-231 cell adhering to a micropillar array. Exactly the same format as Fig. 2 (here Δ*θ* = 180^*°*^).

**Supporting Fig 2.**
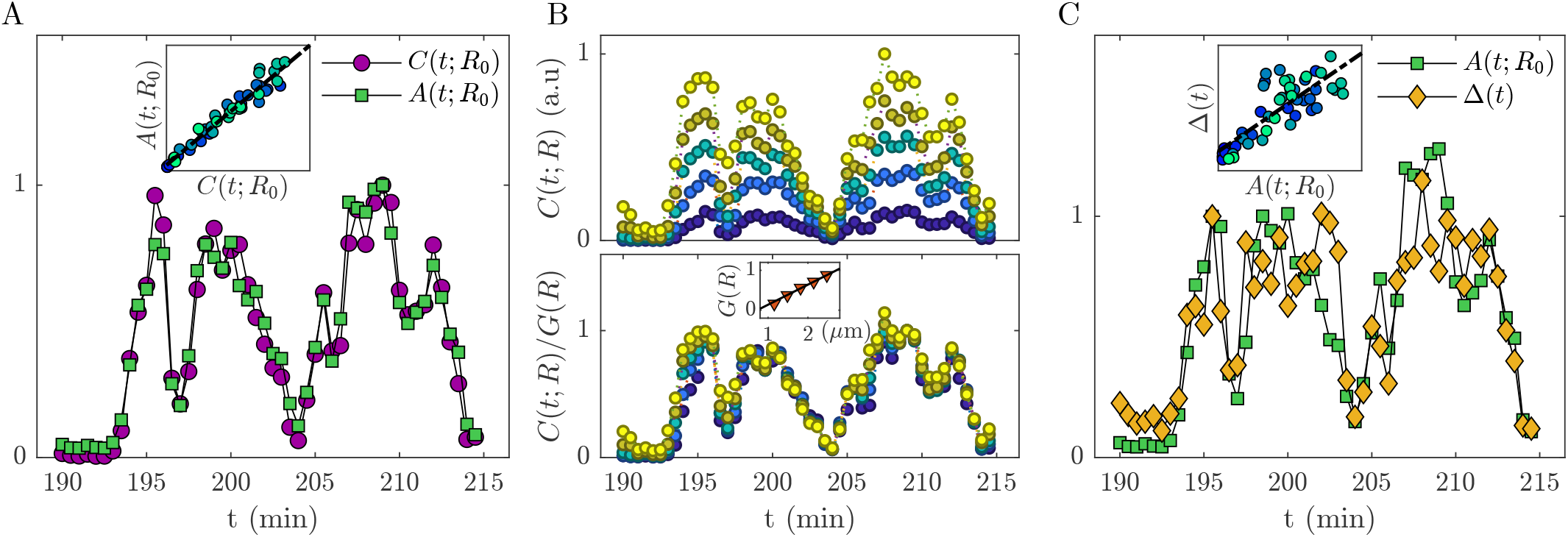
An additional example of an isolated adhesion event in a quasi-stationary spreading MDA-MB-231 cell adhering to a micropillar array. Exactly the same format as Fig. 2 (here Δ*θ* = 180^*°*^). Note that here the adhesion almost disassembles around *t* = 204 min (and likewise the generated force almost drops to zero), yet the adhesion subsequently reassembles (and likewise the force is rebuilt). Despite the different dynamics compared to Fig. 2 and Supporting Fig. 1, Eqs. (1)-(3) (with *G*(*R*) *∼ R*) are remarkably satisfied.

**Supporting Fig 3.**
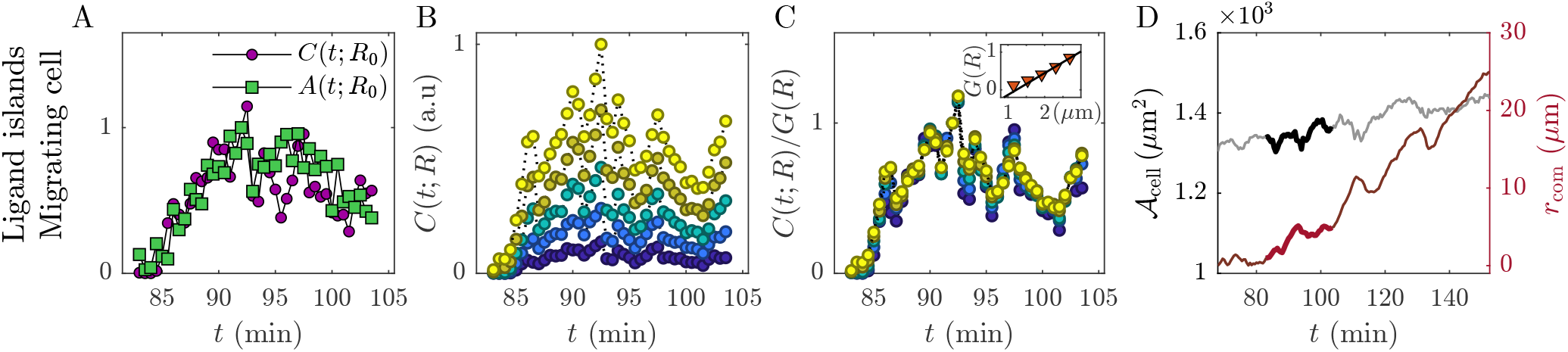
A counterpart of the top row of Fig. 3, presenting an additional example. The integration angular sector here is Δ*θ* = 120^*°*^.

**Supporting Fig 4.**
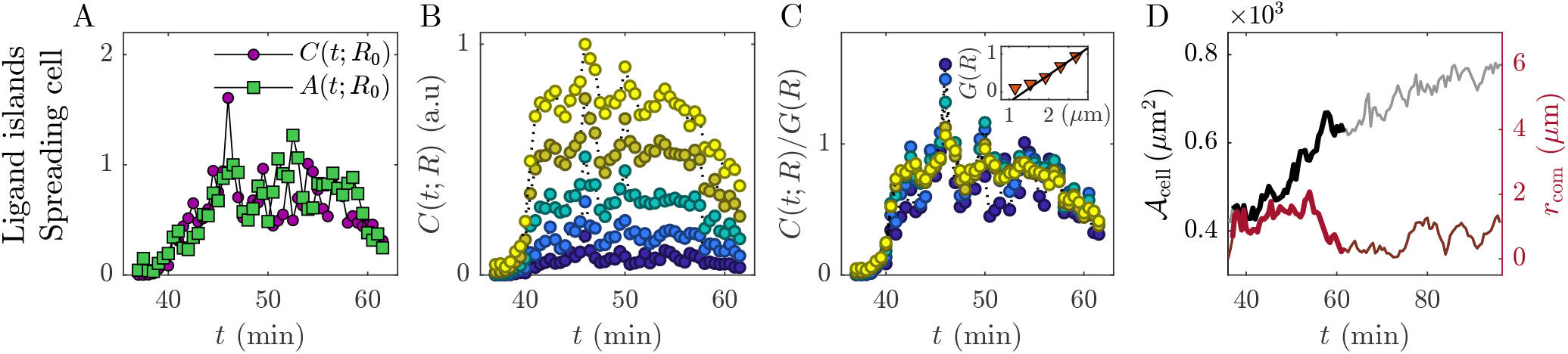
A counterpart of the top row of Fig. 3, presenting an additional example. Note that here, unlike the migratory cell state considered in the top row of Fig. 3 and in Supporting Fig. 3, the cell is actively spreading, see (D). Here we also have Δ*θ* = 135^*°*^.

**Supporting Fig 5.**
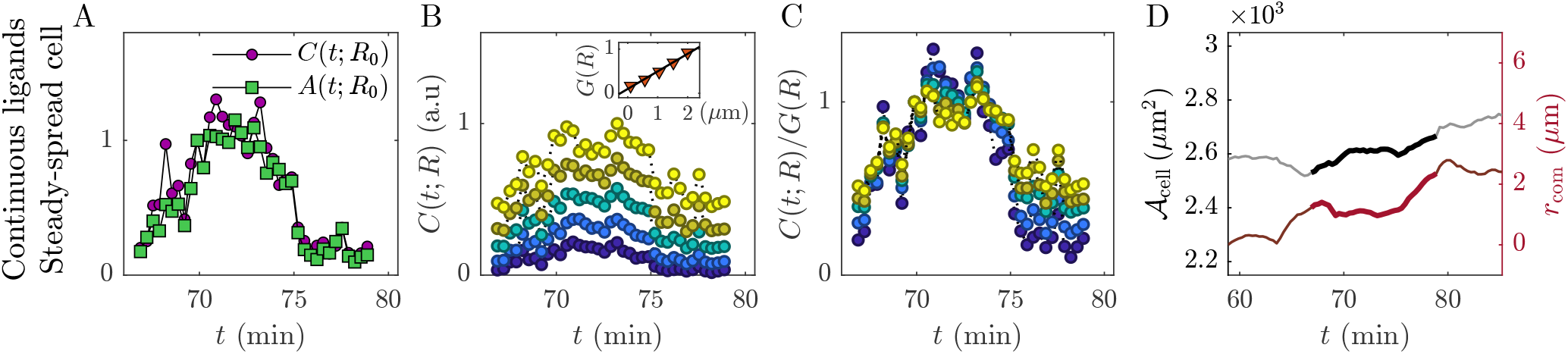
A counterpart of the bottom row of Fig. 3, presenting an additional example. Exactly the same format is used except for the panel labels, which are (E)-(H) in the bottom row of Fig. 3.

**Supporting Fig 6.**
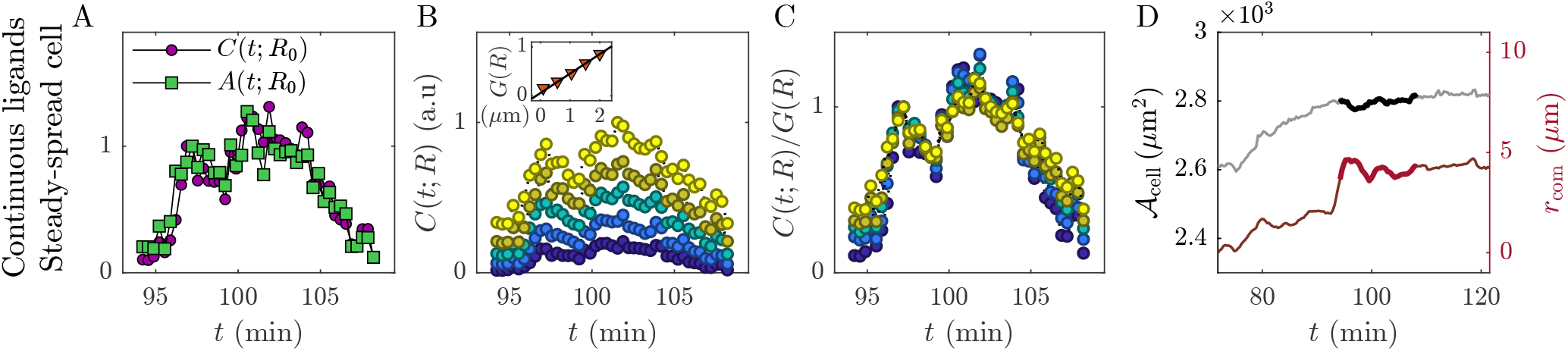
A counterpart of the bottom row of Fig. 3, presenting an additional example. Exactly the same format is used except for the panel labels, which are (E)-(H) in the bottom row of Fig. 3.

## Supplementary Text

The goal of this document is to provide technical details about the results presented in the manuscript, along with some additional supporting results.

### S-1. Identifying “isolated adhesion events”

In this work, we analyzed many “isolated adhesion events” that allow us to rather cleanly associate the time-dependent contractile force with the spatiotemporal amounts of adhesion and F-actin that are involved in generating it. The challenge here is to identify adhesion events in which the potential interference from other, coexisting adhesions in their vicinity is minimal. This is achieved manually (i.e., not automatically) as follows.

First, in both the micropillar array approach and the micropatterned ECM approach, adhesions can form only at well-defined and a priori known potential locations where ligands are available (fibronectin is used through-out this work, see above). Moreover, as we are mostly interested in adhesions that are formed near the cell edge, where the associated generated contractile force is of prime importance (e.g., for cell spreading, cell migration and cellular mechanosensing), the actual number of potential adhesion sites is in fact rather constrained. The temporal onset of a given adhesion event is accurately tracked once the adhesion density field *a*(*r, θ, t*) rises above the background noise level at one of the lig- and islands (i.e., the top surface of a near cell edge pillar in the micropillar array approach or a micropatterned ligand domain in the micropatterned ECM approach). In the former case, a contractile force/displacement Δ(*t*) typically simultaneously emerges.

Second, while tracking the spatiotemporal evolution of *a*(*r, θ, t*) during the adhesion lifetime is straightforward, the challenge is to properly identify the F-actin field *c*(*r, θ, t*) that is associated with a given adhesion, in the sense that it can potentially contribute to the corresponding contractile force being generated. A contractile force, as the term implies, is directed from the adhesion towards the cell body. While this process takes place, i.e., during the lifetime of an adhesion under consideration, the cell edge and the accompanying F-actin network can in fact spread further away and form new adhesion(s) that may coexist with the adhesion of interest. The force generated by such new adhesion(s) in this simultaneous process typically counteracts the contractile force of the adhesion event being analyzed and *c*(*r, θ, t*) that is formed beyond the relevant adhesion cannot possibly contribute to the contractile force of interest.

The spatial separation between adjacent micropillars in the micropillar array approach and the spatial separation between ligand islands in the micropatterned ECM approach tend to separate the two processes (though they might also be naturally separated in cellular dynamics even in the absence of a geometric ligand constraint). Yet, if the two processes are not well separated in time (and hence also not in space), we judge the adhesion event not to be “isolated”, i.e., to experience interference from nearby, coexisting adhesions. The very same biophysical considerations also imply that we should take into account the relevant part of *c*(*r, θ, t*) that is associated with a given adhesion event (and hence with *a*(*r, θ, t*)) and with the corresponding contractile force, which generally points towards the cell body. This gives rise to the integration angular sector Δ*θ* mentioned in the manuscript in relation to the coarse-graining of the basic density fields, to be discussed below (see also movie S1, where the spatiotemporal dynamics of the isolated adhesion event analyzed in Fig. 2 are shown).

In the third experimental approach used in the manuscript, involving a continuous ligand coverage of continuous substrates, the identification of isolated adhesion events is more challenging. Note that the position and bounding region of the adhesions are self-selected by the cell in this case. Yet, the above-stated conditions for the existence of isolated adhesion events can be naturally met also in this experimental approach, and indeed such adhesion events have been carefully identified and analyzed. In practice, we apply the above-discussed criteria based on repeated inspection of time-lapse movies, as also done above, closely following the adhesion density field. The onset of an isolated adhesion event is determined when the latter increases above background noise level near the cell edge and only when no neighboring adhesions emerged in their vicinity throughout their lifetime. This enabled extended tracking over time and across several micrometers surrounding the adhesion center without interference from adjacent adhesion signals. Selected adhesions were tracked from formation to disassembly by following them both forward and backward in time, and adhesion boundaries were determined using TrackMate-Weka (Fiji/ImageJ).

Concrete examples showing spatial snapshots of isolated adhesion events (see Fig. S1), along with a movie that illustrates the spatiotemporal aspects of them (see movie S1), will be provided next.

### S-2. Spatial coarse-graining

In this work, we associate with each isolated adhesion event coarse-grained amounts of F-actin and adhesion (and the contractile force in the micropillar array approach). Here, we provide additional details about how these coarse-grained quantities are obtained from the raw experimental data. The fluorescence microscopy measurements (see Methods), for either an adhesion protein (vinculin or paxillin in our experiments) or F-actin, are performed with a spatial discretization Δ*r* (linear pixel size) and a temporal discretization Δ*t*. These measurements can be then used to construct the adhesion density field *a*(*x, y, t*) and the F-actin density field *c*(*x, y, t*) throughout the entire field of view (see Methods). Here, (*x, y*) corresponds to a planar Cartesian coordinate system, where the integration over the cell depth dimension *z* is automatically performed by the fluorescence intensity measurement (see Methods).

Once an isolated adhesion event is identified, as explained above, these density fields can be more naturally associated with it. In the manuscript, we refer to the micropillar array approach, where the most natural coordinate system is a polar one (*r, θ*) located at the center of the circular top surface of a relevant pillar, see Fig. S1A. Then, as explained in the manuscript, the coarsegrained quantities 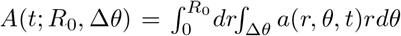 and 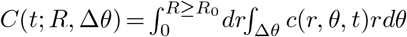 (*R*_0_ is the radius of the top surface of the pillar) are obtained, once the angular sector Δ*θ* is selected. As explained above, Δ*θ* is selected so as to account for the part of *c*(*r, θ, t*) that could possibly contribute to the contractile force. The latter, represented by the vector 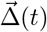 in Fig. S1A (in the manuscript, we report 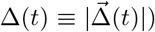), obviously points towards the cell body (i.e., away from the cell edge, see Fig. S1A).

In the micropillar array experiments, as already demonstrated in previous work [9], F-actin and the adhesion itself do not cover the entire ligand-coated top surface of a micropillar but rather form an annulus (see Fig. S1A, apparently due to the non-planar ECM topography, an issue that is not discussed here). Under these conditions, one expects that the part of the annulus that points towards the cell edge would contribute less to contractile force generation compared to the part that points towards the cell body, where the F-actin network is more mature/extended and there are more available sites for myosin-II motors to act. Consequently, we typically selected Δ*θ* to be of a sizable fraction of 360^*°*^ (2*π* radians) and to point towards the proximal part of the adhesion, see Fig. S1A. Yet, since our main findings are in fact valid for small spatial patches and narrow angular windows — as highlighted in the manuscript — we do not expect any significant dependence on the choice of Δ*θ*. This is explicitly demonstrated in Fig. S2A. Finally, note that when we report results for *R* = *R*_0_, the latter is understood to within a single pixel resolution, i.e., we in fact use *R* = *R*_0_ + Δ*r* to ensure that we indeed account for the entire adhesion. The above-described coarse-graining process is explicitly demonstrated in movie S1 for the isolated adhesion event presented in Fig. 2.

In the micropatterned ECM approach, the same considerations discussed above in the context of the the micropillar array approach apply, with one difference: in the former, adhesion proteins cover the circular ligand island more uniformly, i.e., annuli are not formed for planar ECMs, and hence it is natural to define the angular sector Δ*θ* in these experiments with respect to a point on the island circumference close to cell edge, see Fig. S1B. The spatial integration domain is defined in a ‘composite manner’ such that we first consider the area defined by the angular sector Δ*θ* with *r* = *R*_0_ relative to the aforementioned circumferential point and then add to it the area defined by the same Δ*θ* with Δ*r* = *R ≥ R*_0_ relative to the center of the ligand island, see Fig. S1B. That way, the angular sector Δ*θ* for *R* = *R*_0_ is an inscribed angle of the island. The *R* and Δ*θ* notation and values used in the top row of Fig. 3 follow this definition, which is slightly different from the one used in Fig. 2, as explained above. Specifically, we used *R/R*_0_ = 1.15, 1.54, 1.92, 2.30, 2.69 in Fig. 3B (from the bottom curve to the top one) with Δ*θ* = 120^*°*^. The robustness against the specific choice of Δ*θ* in the micropatterned ECM approach is explicitly demonstrated in Fig. S2B. The results in Fig. S2 in which *C*(*t*; *R*, Δ*θ*) for different values of Δ*θ* is shown to approximately collapse on a master curve once normalized by its value at a specific point in time, denoted as *F* (Δ*θ*), imply that *G*(*R*; Δ*θ*) — discussed in the manuscript and in Sect. S-3 below — might be further decomposed multiplicatively as *G*(*R*; Δ*θ*) = *G*(*R*)*F* (Δ*θ*). These findings further reinforce the local nature of our main results, which underlie the relations we observed between the various coarse-grained quantities.

In the experimental approach that involves a continuous ligand coverage of planar, continuous ECMs, the integration domain used for coarse-graining is defined as follows; at any time *t*, we identify the area that corresponds to the isolated adhesion event, enclosed by the blue line and highlighted in white in Fig. S1C. This area (and its shape), which is self-selected by the cell, constitutes the analog of the fixed circle of radius *R*_0_ in the two other experimental approaches and symbolically corresponds to *R* = *R*_0_ in the bottom row of Fig. 3. Likewise, we denote by *R ≥ R*_0_ an integration domain that is obtained by a self-similar expansion by a factor *R/R*_0_ *≥* 1 of the enclosing blue line, see the yellow line in Fig. S1C.

All the above-discussed coarse-graining involved areal integration. At the bottom of Fig. 4A, we also present coarse-grained quantities obtained through line integration. This is achieved in two steps. First, a kymograph (i.e., a space-time plot) is generated by selecting a line in space (in our case the line goes through the adhesion, as shown in the top part of Fig. 4D-E), see movie S2. The fluorescent intensity of some label is then plotted at each time snapshot as a function of the spatial coordinate along the line. As the line has a finite thickness (of a few pixels), the intensity is integrated over the thickness. Several resulting kymographs are presented in Fig. 4, where we extensively discuss the role of retrograde F-actin flow in contractile force generation. Second, denoting the orientation of the line by *θ*_*i*_ and the coordinate along it by *r*_*i*_, the results at the bottom of Fig. 4A are obtained through a line integral over *r*_*i*_, where the outcome is schematically denoted as *A*(*t*; *R*_0_, *θ*_*i*_) and *C*(*t*; *R*_0_, *θ*_*i*_) (see legend therein).

**Fig S1.**
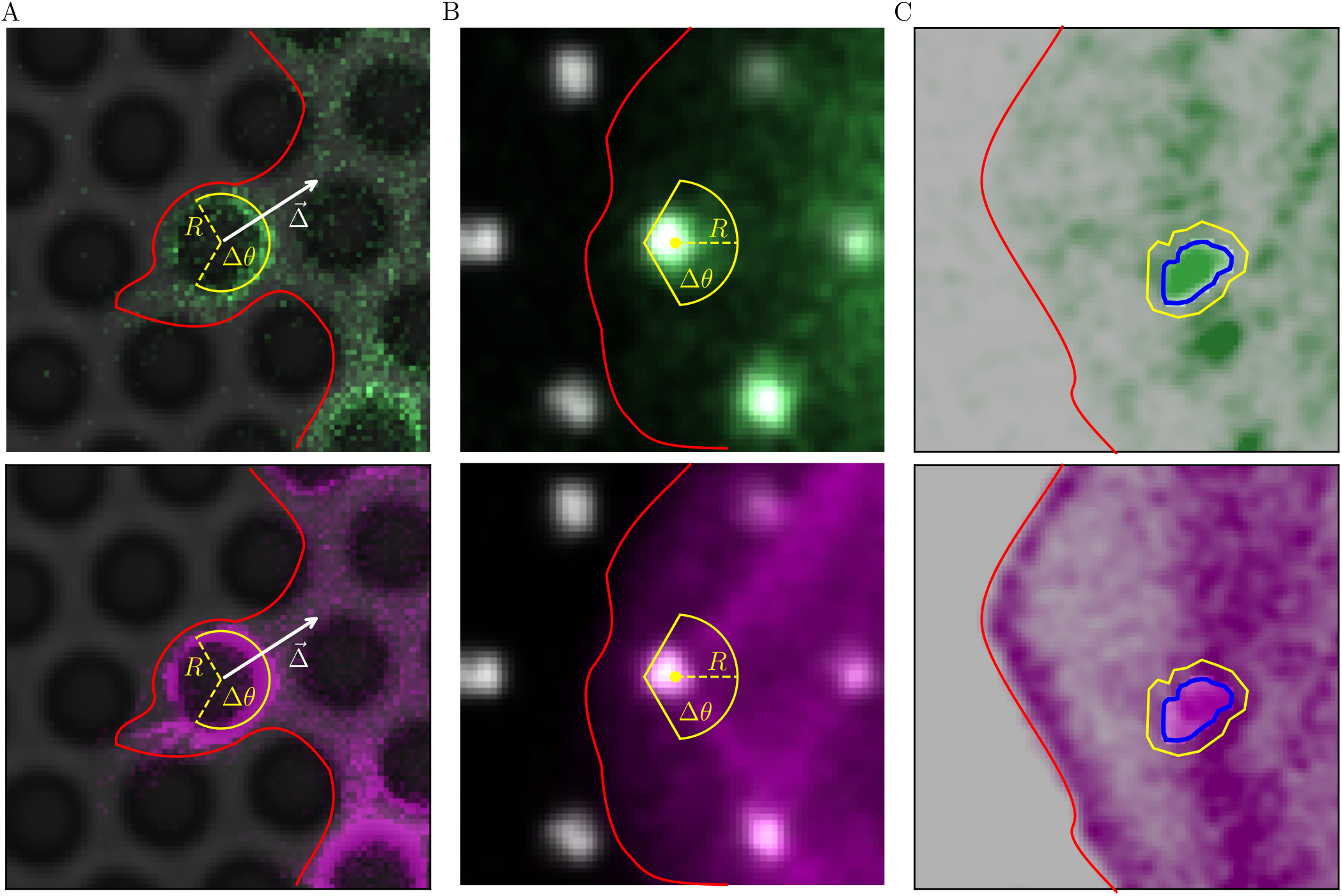
Spatial coarse-graining in the three experimental approaches. In each column, the same time snapshot in a given experiment is presented twice (zoomed-in on part of the cell edge), one showing F-actin (green, top) and one an adhesion protein (magenta, bottom). The red line marks the cell edge location. (**A**) Snapshots of an isolated adhesion event in a micropillar array experiment, corresponding to *t* = 226.5 min in Fig. 2. The top surface of each pillar appears in black, even though it is actually coated by fibronectin (not imaged here). The adhesion protein (here vinculin) does not uniformly covers the fibronectin-coated top surface of the pillar and the same applies to F-actin, as shown in previous work [9]. The angular sector Δ*θ* is marked. Together with *r* = *R*, they form the spatial integration domain used for coarse-graining, see text. The contractile displacement vector 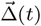 at this snapshot is marked as well (its magnitude is arbitrary, chosen for visual clarity). (**B**) The same as (A) but for an isolated adhesion event in an experiment employing a micropatterned ECM with a hexagonal array of circular fibronectin islands (white circles), corresponding to *t* = 96.5 min in the top row of Fig. 3. Here, the adhesion protein is paxillin. The the spatial integration domain used for coarse-graining is marked, see text for extended discussion (and for the meaning of the angular sector Δ*θ* and *R* here). (**C**) The same as (A) but for an isolated adhesion event in an experiment employing a continuously fibronectin-coated ECM, corresponding to *t* = 71.55 min in the bottom row of Fig. 3. Here, the adhesion protein is paxillin. See text for the identification of the adhesion in this case and for the definition of the spatial integration domain used for coarse-graining.

### S-3. The near-adhesion, spatial distribution of F-actin

*∼*

A major result in the manuscript is the multiplicative space-time decomposition of the coarse-grained F-actin amount *C*(*t*; *R*, Δ*θ*) in Eq. (2), i.e., *C*(*t*; *R*, Δ*θ*) = *G*(*R*, Δ*θ*) *T* (*t*), where we find *G*(*R*, Δ*θ*) *∼ R* to a good approximation. It is further stated that the latter relations indicate that the coarse-grained F-actin density follows *c*(*r, θ, t*) *∼ g*(*r*)*T* (*t*) with *g*(*r*) *∼* 1*/r*, where 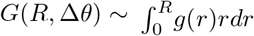. Here, we further substantiate these basic properties of the near-adhesion, spatial distribution of F-actin and their implications. In Fig. S3A, we present *G*(*R*, Δ*θ*) of the inset of Fig. 2B(bottom) (with the same shorthand notation *G*(*R*)). Recall that *G*(*R*) is the amplitude of *C*(*t*; *R*, Δ*θ*) in Fig. 2B(top) at a given snapshot, i.e., its value at a fixed time *t. G*(*R*) is defined and extracted in this way throughout this work. It is observed that at distances further from the edge of the adhesion (located at *r* = *R*_0_ = 1 *µ*m), *G*(*R*) *∼ R* to a good approximation (indicated by the superposed dashed line).

Using the above relations and the predominantly linear variation of *G*(*R*) with *R*, the relation 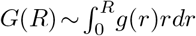 indicates that *g*(*r*) *∼* 1*/r*, as stressed above and in the manuscript. We independently tested the latter relation in Fig. S3B, where we present *g*(*r*) as obtained by an angular average of the measured *c*(*r, θ, t*) (i.e., *θ*-average over Δ*θ*), for the same time *t* considered in Fig. S3A. Indeed, we find *g*(*r*) *∼* 1*/r* slightly away from the adhesion edge (indicated by the superposed dashed line). This inhomogeneous, radial decay of the F-actin density *g*(*r*) is generically observed in our experiments. It appears to be qualitatively consistent with the measurements of [51], see in particular Fig. 3D therein. Moreover, and importantly, a *g*(*r*) *∼* 1*/r* decay of the coarse-grained F-actin density (corresponding to a constant amount of F-actin in each shell of fixed thickness *dr*) is consistent with a radial growth of quasi-one-dimensional (linear) actin filaments from small adhesion patches, as highlighted in the manuscript.

**Fig S2.**
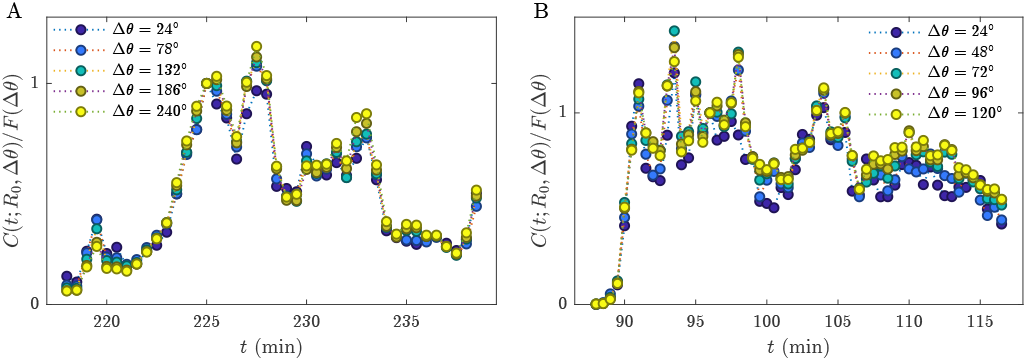
Robustness against the specific choice of a spatial integration angular sector Δ*θ*. (**A**) Similarly to the analysis in Fig. 2B but for *C*(*t*; *R*_0_, Δ*θ*), i.e., a fixed *R* = *R*_0_ and various choices of Δ*θ* (see legend, where Δ*θ* = 240^*°*^ has been used in the manuscript, see also Fig. S1A). The function *F* (Δ*θ*) used to normalize *C*(*t*; *R*_0_, Δ*θ*) (see y-axis label) corresponds to its value at a specific point in time. (**B**) The same as (A) but similarly to the analysis in Fig. 3C (see legend for the values of Δ*θ* and note that Δ*θ* = 120^*°*^ has been used in the manuscript, see also Fig. S1B). The robustness of the results against the specific choice of Δ*θ* is demonstrated, see text for additional discussion.

**Fig S3.**
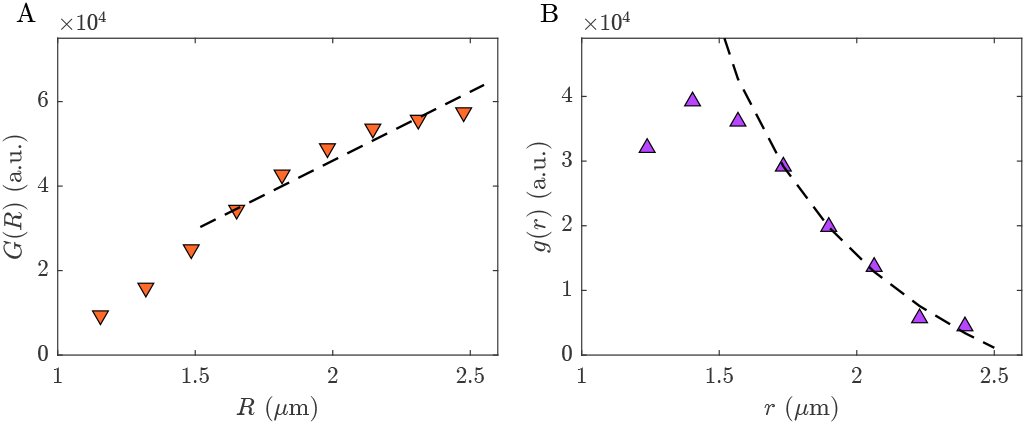
The near-adhesion, spatial distribution of F-actin as quantified by the functions *G*(*R*) **and** *g*(*r*). (**A**) *G*(*R*, Δ*θ*) of the inset of Fig. 2B(bottom) (with the same shorthand notation *G*(*R*)), here presented at a higher spatial resolution for enhanced visual clarity. The superposed dashed line highlights the predominantly linear variation of *G*(*R*) with *R* slightly away from the edge of the adhesion at *R*_0_ = 1 *µ*m. (**B**) The independently computed *g*(*r*) as obtained by an angular average of the measured *c*(*r, θ, t*) (i.e., *θ*-average over Δ*θ*), for the same time *t* considered in (A). The expected *g*(*r*) *∼* 1*/r* behavior slightly away from the adhesion edge is indicated by the dashed line. See text for discussion.

### S-4. The robustness and generality of the experimental observations

In Figs. 2-3, we analyzed in detail one isolated adhesion event for each of the three distinct experimental approaches employed in this work, supporting the validity of Eqs. (1)-3) therein with *G*(*R*) *∼ R*. It is also stated in the manuscript that the experimental observations are in fact valid for many other adhesion events in three experimental approaches. We presented additional examples in Supporting Figs. 1-6 to support this statement. Many more examples are provided in [52].

### S-5. Supplementary movies

This manuscript contains two supplementary movies, described as follows:

#### Movie S1

The movie shows part of the edge of an MDA-MB-231 cell co-expressing GFP-vinculin (green) and tdTomato-Tractin (magenta) spreading on a fibronectin-coated micropillar array (the same one shown in Fig. 2). A single adhesion is shown which was manually tracked from formation to disassembly within the marked angular sector Δ*θ*. The associated F-actin accumulation and pillar displacement were quantified in parallel. The plots on the right show the temporal evolution of adhesion intensity, F-actin intensity, and pillar displacement-derived force generation.

#### Movie S2

Themovie shows part of the edge of a MEF (the same one shown in Fig. 4A) co-expressing GFP-paxillin (green) and tdTomato-Tractin (magenta) spreading on a fibronectin micropatterned substrate (fibronectin pattern not shown). To capture rapid actin dynamics, F-actin images were acquired in bursts of 20 consecutive frames between paxillin images collected at 30-s intervals. The red rectangle indicates the region used to generate the kymograph shown in Fig. 4A.

